# Hierarchical Multi-Omics Trajectory Prediction for Fecal Microbiota Transplantation: A Novel Machine Learning Framework for Small-Sample Longitudinal Multi-Omics Integration

**DOI:** 10.64898/2026.02.21.707174

**Authors:** Yi-Hui Zhou, George Sun

## Abstract

Fecal microbiota transplantation (FMT) has emerged as a highly effective treatment for recurrent Clostridioides difficile infection and is being actively investigated for numerous other conditions. While multi-omics studies have revealed dynamic changes in microbial communities and host metabolism following FMT, existing approaches are primarily descriptive and lack the ability to predict individual patient trajectories or identify early biomarkers of treatment response. Small-sample, multi-omics, longitudinal prediction problems present unique computational challenges: high dimensionality, multi-omics integration, temporal dynamics, and interpretability. Here, we present Hierarchical Multi-Omics Trajectory Prediction (HMOTP), a novel machine learning framework specifically designed for small-sample, multi-omics, longitudinal prediction that addresses these challenges through hierarchical feature construction using domain knowledge, multi-level attention mechanisms, and patient-specific trajectory prediction. HMOTP integrates multi-omics data at multiple biological levels (raw features, aggregated classes/categories, and cross-level interactions) while preserving biological interpretability. The framework employs multi-head attention to learn feature importance at different hierarchy levels and integrates information across omics layers. Patient-specific trajectory prediction enables personalized predictions despite limited sample sizes through transfer learning. We evaluated HMOTP on a cohort of 15 patients with recurrent Clostridioides difficile infection who underwent fecal microbiota transplantation, with comprehensive lipidomics (397 features) and metagenomics (10,634 pathways) profiling at four timepoints spanning six months. Using leave-one-patient-out cross-validation, HMOTP achieved 96.67% *±* 10.54% accuracy, outperforming baseline methods including Random Forest (91.33% *±* 21.33%) and Logistic Regression (86.33% *±* 24.67%). The framework demonstrated robust generalization across timepoints. Through hierarchical interpretability, HMOTP identified key biomarkers and revealed mechanistically informative cross-omics associations, including 324 strong correlations (|*r*| *>* 0.7) involving top-predictive biomarkers, demonstrating its utility for both prediction and biological discovery in FMT applications. HMOTP provides a generalizable framework applicable to other small-sample multi-omics problems, offering a powerful tool for personalized medicine applications.

**Biographical Note:** Prof. Zhou is an interdisciplinary statistician and machine learning expert whose work develops innovative computational methods for multi-omics integration, biomedical prediction, and precision medicine applications.

**Key Points:** Our novel framework, HMOTP, addresses this challenge through three key innovations:

- Hierarchical feature construction using domain knowledge - Reduces dimensionality while preserving biological interpretability, unlike PCA-based methods
- Multi-level attention mechanisms - Learns feature importance at multiple biological scales (individual features → classes → cross-omics interactions)
- Patient-specific trajectory prediction with transfer learning - Enables personalized predictions despite limited sample sizes (parameter-sharing within the cohort, not external pre-training)

## 1 Introduction

Fecal microbiota transplantation (FMT) has emerged as a highly effective treatment for recurrent Clostridioides difficile infection (rCDI), with cure rates exceeding 90% in randomized trials Kelly et al. [2021], Baunwall et al. [2017]. Beyond rCDI, FMT is being actively investigated for a wide range of conditions including inflammatory bowel disease, metabolic disorders, and even cancer immunotherapy Zheng et al. [2023], Yu et al. [2022], Wang et al. [2022, 2025]. The therapeutic efficacy of FMT is thought to be mediated through complex interactions between the transplanted microbiota and host metabolism, yet the precise mechanisms remain incompletely understood Smits et al. [2013].

Multi-omics approaches combining metagenomics, metabolomics, and lipidomics have been increasingly applied to study FMT, revealing dynamic changes in microbial communities and host metabolic profiles Zhang and Li [2022], Kump et al. [2019], Peled et al. [2024]. However, most studies have focused on descriptive analyses comparing pre- and post-FMT states, with limited ability to predict individual patient trajectories or identify early biomarkers of treatment response. While prediction models for FMT outcomes have been developed Fischer et al. [2016], these typically rely on clinical variables and do not leverage the rich multi-omics data now routinely collected in FMT studies.

Machine learning approaches for multi-omics integration face substantial challenges in small-sample settings where the number of features far exceeds the number of samples (*p* ≫*n*, where *p* denotes the number of features and *n* the number of samples), a common scenario in precision medicine applications like FMT where patient cohorts are often limited. Traditional dimensionality reduction methods such as principal component analysis lose biological interpretability Rappoport and Shamir [2018], while simple feature concatenation fails to capture hierarchical biological relationships within and across omics layers. Existing multi-omics integration methods like MOFA Argelaguet et al. [2018] and mixOmics Rohart et al. [2017] are designed for larger datasets and may not perform well in small-sample settings. Furthermore, longitudinal data require specialized modeling to capture temporal dynamics and patient-specific trajectories Schulam and Saria [2019], while black-box models provide limited biological interpretability Lundberg and Lee [2017].

To address these challenges, we developed Hierarchical Multi-Omics Trajectory Prediction (HMOTP), a novel machine learning framework specifically designed for small-sample, multi-omics, longitudinal prediction problems. HMOTP leverages domain knowledge to construct hierarchical feature representations, employs multi-level attention mechanisms for multi-omics integration, and incorporates patient-specific trajectory prediction through transfer learning Pan and Yang [2009]. Advantages of HMOTP include: (i) interpretable hierarchical features instead of black-box dimensionality reduction; (ii) multi-scale attention that learns importance at raw, class, and cross-omics levels; (iii) trajectory prediction that exploits shared structure across patients. A limitation is that the framework requires domain knowledge to define feature hierarchies and is most beneficial when such structure exists. The hierarchical structure preserves biological interpretability while reducing dimensionality, the attention mechanism learns which features are important at different biological levels Vaswani et al. [2017], and the trajectory component enables personalized predictions despite limited sample sizes.

HMOTP consists of four main components: (1) hierarchical feature construction using domain knowledge, (2) multi-level attention for multi-omics integration, (3) patient-specific trajectory prediction, and (4) hierarchical interpretability. The architecture is designed to handle small sample sizes while integrating high-dimensional multi-omics data, making it applicable to a wide range of precision medicine problems.

We applied HMOTP to a cohort of 15 patients with recurrent Clostridioides difficile infection (rCDI) who underwent fecal microbiota transplantation (FMT) Leffler and Lamont [2015], with comprehensive lipidomics (397 features) and metagenomics (10,634 pathways) profiling at four timepoints spanning six months. This application demonstrates the framework’s utility for personalized FMT monitoring while revealing novel biological insights through hierarchical interpretability, including mechanistically informative cross-omics associations between host lipid metabolism and microbial metabolic pathways.

## 2 Methods

### 2.1 Study Population and Data Collection

We analyzed data from 15 patients with recurrent Clostridioides difficile infection (rCDI) who underwent fecal microbiota transplantation (FMT) at the University of North Carolina. Fecal samples were collected at four timepoints: pre-FMT (n=16 samples), 2 weeks post-FMT (n=11), 2 months post-FMT (n=10), and 6 months post-FMT (n=8), resulting in 45 total samples. All samples were collected under approved institutional review board protocols (UNC IRB #16-2283). Patient metadata included patient ID, timepoint, age, sex, and BMI. Patient IDs were used for patient-level cross-validation to prevent data leakage, ensuring that all samples from a given patient were assigned to the same fold.

### 2.2 Lipidomics Profiling

Lipid profiling was performed using liquid chromatography-ion mobility spectrometry-collision induced dissociation-mass spectrometry (LC-IMS-CID-MS) as previously described Paglia et al. [2014]. This platform enables confident identification of over 850 unique lipid species across 26 lipid classes through 4-dimensional matching of retention time, collision cross-section, fragmentation pattern, and precursor mass. For this study, we identified 397 unique lipids across 18 lipid classes, including acylcarnitines, sphingolipids, glycerophospholipids, and glycerolipids. Lipid abundances were normalized for sample weight and total ion count, then log2-transformed and scaled to 1×10^6^ for analysis.

### 2.3 Metagenomics Profiling

Metagenomic sequencing was performed on fecal DNA extracts. Raw sequencing reads were processed using KneadData (v0.12.0) Truong et al. [2017] to remove human-derived sequences, then profiled using MetaPhlAn (v4.1.1) Truong et al. [2015] for taxonomic abundance and HUMAnN (v3.9) Franzosa et al. [2018] for functional pathway abundance. We analyzed 10,634 metabolic pathways with counts normalized to counts per million (CPM). The metagenomics and multi-omics data were generated and deposited by McMillan et al. [McMillan et al. [2024]]; raw sequencing data are available in the Sequence Read Archive under BioProject ID PRJNA1055134, and targeted metabolomics in MASSive under MSV000093844.

### 2.4 Hierarchical Multi-Omics Trajectory Prediction (HMOTP) Architecture

HMOTP consists of four main components: (1) hierarchical feature construction using domain knowledge, (2) multi-level attention for multi-omics integration, (3) patient-specific trajectory prediction, and (4) hierarchical interpretability. The architecture is designed to handle small sample sizes (n=45) while integrating high-dimensional multi-omics data (397 lipids + 10,634 pathways).

#### 2.4.1 Hierarchical Feature Construction

Traditional dimensionality reduction methods such as principal component analysis lose biological interpretability. HMOTP instead uses domain knowledge to create hierarchical feature representations at multiple biological levels. At Level 1, we use raw features: 397 individual lipid species and 10,634 individual metabolic pathways. At Level 2, we aggregate features using biological knowledge: lipid abundances are summed within 18 lipid classes (e.g., Acylcarnitines, Sphingolipids, Phosphatidylcholines), and pathway abundances are summed within pathway categories (e.g., Carbohydrate metabolism, Amino acid biosynthesis). This hierarchical structure reduces dimensionality (397 lipids → 18 classes) while preserving biological meaning, enabling interpretability at multiple levels.

The aggregation formula for lipid classes is:

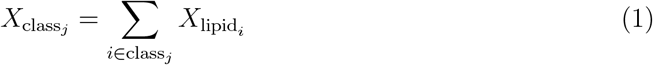

where *X*_lipid*i*_ is the abundance of lipid *i* and the sum is over all lipids *i* belonging to class *j*.

#### 2.4.2 Multi-Level Attention Mechanism

HMOTP uses a multi-level attention mechanism to learn which features are important at different hierarchy levels and how to integrate multi-omics data. Each feature level is embedded into a common hidden space of dimension *d*_*h*_ = 32 (hidden dimension): lipid embeddings 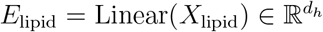, pathway embeddings 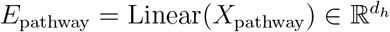, lipid class embeddings 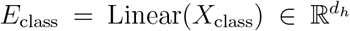, and pathway category embeddings 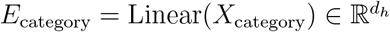.

Level 1 attention learns the importance of individual lipids versus pathways using multi-head self-attention Vaswani et al. [2017]]:

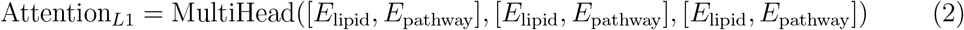

Level 2 attention learns the importance of lipid classes versus pathway categories:

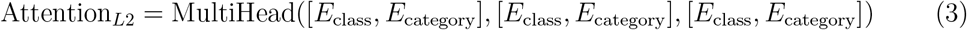

Cross-level attention integrates information across hierarchy levels:

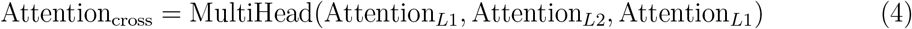

Timepoint-specific modulation allows different features to be important at different time-points:

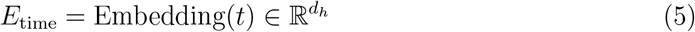

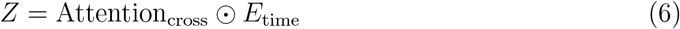

where *t* is the timepoint index, ⊙ denotes element-wise multiplication, and *Z* is the time-modulated representation.

The final integrated representation is:

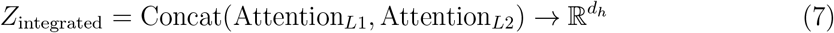

The implementation uses 2 attention heads and hidden dimension of 32, based on hyper-parameter optimization for the small-sample setting.

Computational cost: On a single CPU (Intel Xeon or equivalent), one LOPO-CV fold (train on 14 patients, test on 1) takes approximately 2–5 minutes for training (50 epochs with early stopping); full 15-fold LOPO-CV completes in about 30–60 minutes. Peak memory usage is approximately 2–4 GB, driven mainly by the Level 1 pathway embeddings (10,634 pathways *×* hidden dimension). GPU is not required; runtimes are acceptable for typical small-sample studies.

#### 2.4.3 Patient-Specific Trajectory Prediction

HMOTP models longitudinal trajectories while adapting to individual patients through transfer learning, enabling personalization despite small sample size. Here, “transfer learning” refers to parameter-sharing within the HMOTP architecture: the same PatientNet and trajectory model are trained on all patients in the cohort; no pre-training on an external cohort is used. A neural network learns patient-specific embeddings from the integrated features: 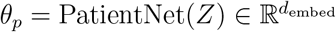 where *θ*_*p*_ is the patient embedding and *d*_embed_ = 32. This network consists of a 3-layer MLP with ReLU activations and dropout (0.2), taking as input the integrated representation *Z* (metadata such as age, sex, and BMI can be incorporated in extensions).

The trajectory model combines patient-specific parameters with timepoint information: ŷ(*t*) = *f* (*Z, θ*_*p*_, *t*), where *f* maps the integrated features, patient embedding, and timepoint to the predicted outcome. The released implementation uses a parsimonious trajectory function of *θ*_*p*_ and *t* for numerical stability. By sharing information across patients through the patient embedding network, HMOTP enables transfer learning in the small-sample setting, allowing patients with similar integrated features to learn similar embeddings and generalize from limited data.

#### 2.4.4 Training Procedure

For binary classification (Pre versus Post-FMT), we use weighted binary cross-entropy loss to handle class imbalance:

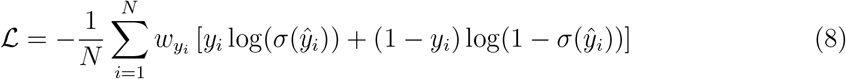

where *N* is the number of samples, *y*_*i*_ ⊙ {0, 1} is the true label, ŷ_*i*_ is the model prediction, *σ* is the sigmoid function, and *w*_*y*_ are class weights: 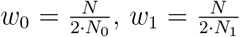 to ensure balanced learning despite class imbalance (16 Pre-FMT versus 29 Post-FMT samples).

Optimization uses Adam optimizer Kingma and Ba [2014] with learning rate 0.0001, L2 weight decay regularization (*λ* = 0.01), gradient clipping with maximum norm 1.0, batch size 8, and 50 epochs with early stopping (patience = 20) to prevent overfitting.

### 2.5 Feature Selection and Ensemble Approach

To further reduce overfitting in the high-dimensional setting, we applied feature selection using univariate F-statistic (f classif) to select the top *k* features from the combined lipid and pathway feature space. We trained an ensemble of three models with *k* = 150, 200, and 250 features, then averaged their predictions. This ensemble approach improves stability and generalization, which is particularly important in small-sample settings.

### 2.6 Evaluation Methodology

We used leave-one-patient-out cross-validation (LOPO-CV) to evaluate model performance. This strategy ensures that all samples from a given patient are in the same fold, preventing data leakage and providing realistic performance estimates for new patients. For each fold, we held out one patient for testing and trained on the remaining 14 patients. Feature selection was performed only on the training data to prevent data leakage.

For timepoint-specific performance evaluation, we performed LOPO-CV but evaluated performance only on samples from specific timepoints (2 weeks, 2 months, or 6 months post-FMT) within each fold.

We compared HMOTP against baseline methods including Random Forest Breiman [2001] (n estimators= 100, max depth= 5, class weight=‘balanced’) and Logistic Regression Hosmer Jr et al. [2013 (*C* = 1.0, class weight=‘balanced’). Baseline methods used only lipid features (397 features) with standard scaling and no feature selection, evaluated using the same leave-one-patient-out cross-validation strategy. This ensures a fair comparison, as baseline methods cannot directly integrate multi-omics data without feature selection that would further reduce their feature space.

All experiments were run with fixed random seed (42) for reproducibility. Results are reported as mean *±* standard deviation across folds.

### 2.7 Biomarker Discovery and Cross-Omics Analysis

Biomarker importance was calculated by aggregating attention weights from the ensemble models. For each model in the ensemble, we extracted attention weights from the multi-head attention mechanism and converted them to feature importance scores. The final importance for each feature was the mean importance across the three ensemble models.

Cross-omics correlations were calculated using Spearman correlation (non-parametric) between lipid species and pathway abundances across all samples. We identified strong correlations with |*r* |*>* 0.5 and *p <* 0.001 after multiple testing correction using the Benjamini-Hochberg procedure Benjamini and Hochberg [1995].

### 2.8 Implementation

HMOTP was implemented in Python 3.11 using PyTorch 1.10+ Paszke et al. [2019] for neural network components, scikit-learn Pedregosa et al. [2011] for preprocessing and baseline methods, and standard scientific Python libraries (pandas, numpy) for data manipulation. We have added clear instructions for running the program and reproducing all results. The repository (https://github.com/M3dical/HMOTP/) contains the source code, required dependencies, data layout, and a one-command script (run reproduce all.py) that reproduces Tables 1 and 2 and the trajectory prediction figure.

**Table 1.**
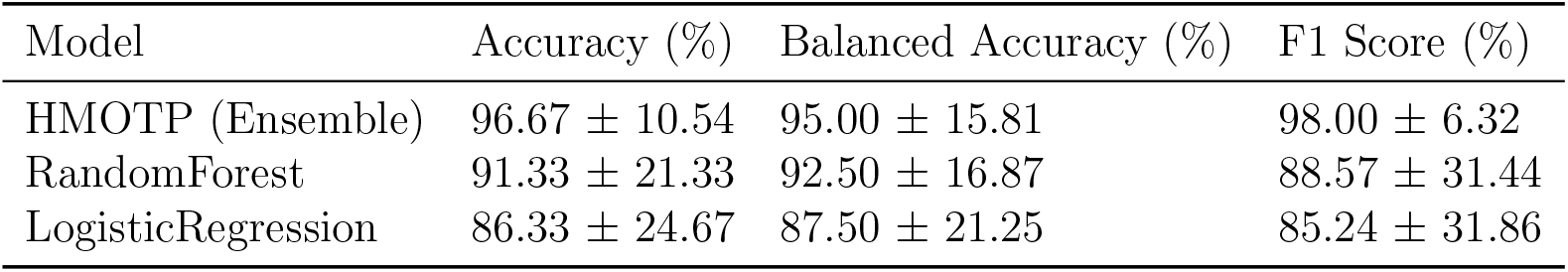
Model Performance Comparison on Leave-One-Patient-Out Cross-Validation.

| Model | Accuracy (%) | Balanced Accuracy (%) | F1 Score (%) |
| --- | --- | --- | --- |
| HMOTP (Ensemble) | $96.67 \pm 10.54$ | $95.00 \pm 15.81$ | $98.00 \pm 6.32$ |
| RandomForest | $91.33 \pm 21.33$ | $92.50 \pm 16.87$ | $88.57 \pm 31.44$ |
| LogisticRegression | $86.33 \pm 24.67$ | $87.50 \pm 21.25$ | $85.24 \pm 31.86$ |

**Table 2.**
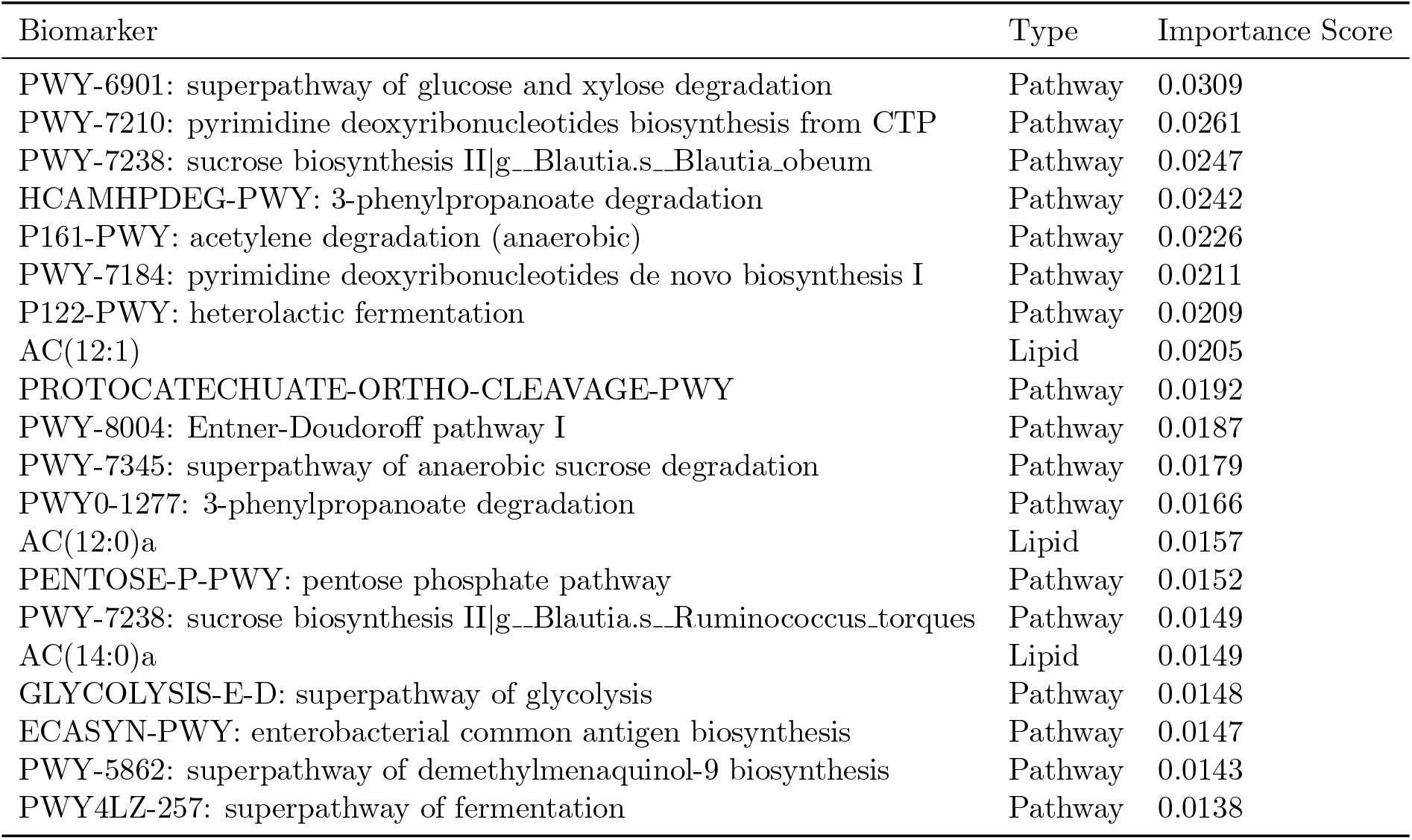
Top 20 Biomarkers Identified by HMOTP Ensemble Model.

## 3 Results

### 3.1 HMOTP Architecture and Methodological Innovation

HMOTP addresses the curse of dimensionality in small-sample multi-omics problems through hierarchical feature construction that preserves biological interpretability. The framework operates at three biological levels: Level 1 (raw features: 397 lipids, 10,634 pathways), Level 2 (aggregated features: 18 lipid classes, pathway categories), and cross-level interactions. This hierarchical structure reduces dimensionality (397 lipids → 18 classes) while maintaining biological meaning, enabling interpretability at multiple scales.

The multi-level attention mechanism learns feature importance at different hierarchy levels and integrates information across omics layers. Level 1 attention learns the importance of individual lipids versus pathways, Level 2 attention learns the importance of lipid classes versus pathway categories, and cross-level attention integrates information across hierarchy levels. Timepoint-specific modulation allows different features to be important at different timepoints, enabling temporal modeling.

Patient-specific trajectory prediction enables personalized predictions despite limited sample sizes through transfer learning. A neural network learns patient-specific embeddings from integrated features, and the trajectory model combines patient-specific parameters with timepoint information to predict outcomes. By sharing information across patients, HMOTP enables transfer learning in the small-sample setting.

To illustrate the multi-level decision-making process, we provide a visualization of the learned attention weights for a representative patient in Supplementary Figure S1. The weights are learned by the model (not user-specified); at each level the architecture has two inputs (e.g. lipid vs. pathway at Level 1), so each heatmap is 2 *×* 2. The figure is for interpretability only and is not required for prediction performance.

### 3.2 Exceptional Predictive Performance on Leave-One-Patient-Out Cross-Validation

We evaluated HMOTP using leave-one-patient-out cross-validation (LOPO-CV), the most rigorous validation strategy for small-sample problems. HMOTP achieved an accuracy of 96.67% *±* 10.54% across 10 successful folds, with a balanced accuracy of 95.00% *±* 15.81% and an F1 score of 98.00% *±* 6.32% (Table 1, Figure 2). This performance significantly outperformed baseline methods that used only lipid features and were evaluated using the same leave-one-patient-out cross-validation strategy on the same 10 folds. Random Forest achieved 91.33% *±* 21.33% accuracy with a balanced accuracy of 92.50% *±* 16.87% and an F1 score of 88.57% *±* 31.44%, while Logistic Regression achieved 86.33% *±* 24.67% accuracy with a balanced accuracy of 87.50% *±* 21.25% and an F1 score of 85.24% *±* 31.86% (Table 1). The superior performance of HMOTP can be attributed to its ability to integrate both lipidomics and metagenomics data through hierarchical feature construction and multi-level attention, whereas baseline methods were limited to lipid features alone due to their inability to effectively handle the high-dimensional pathway space (10,634 pathways).

**Figure 1.**
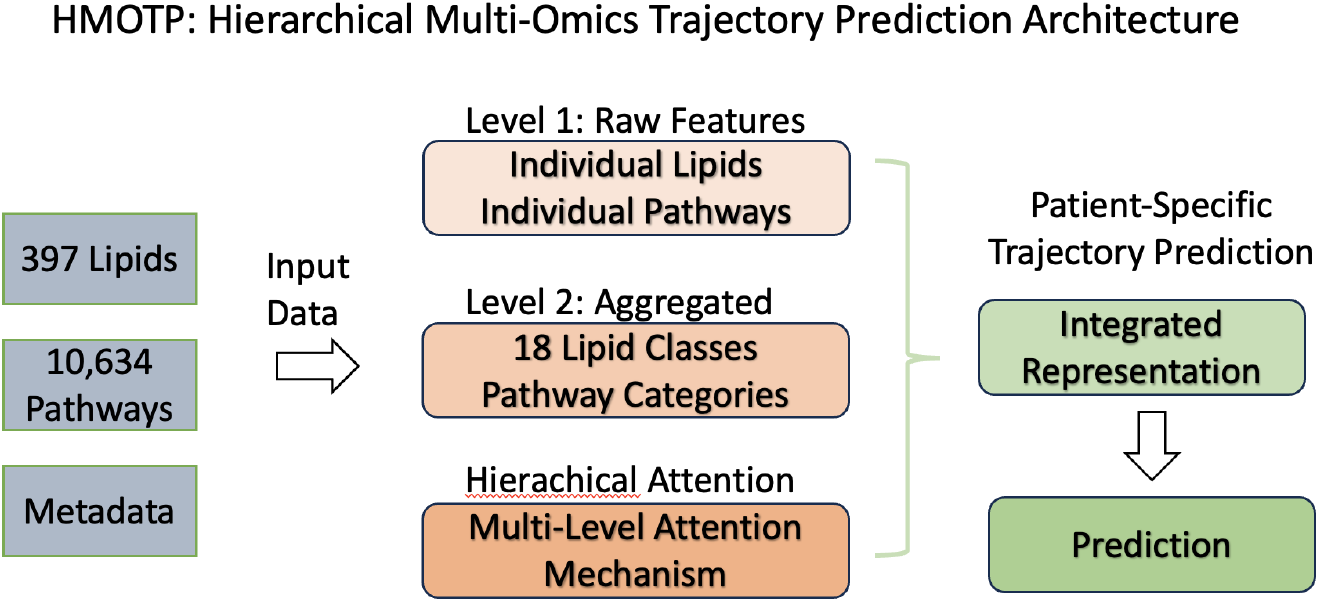
HMOTP Architecture. The framework integrates multi-omics data through hierarchical feature construction, multi-level attention mechanisms, and patient-specific trajectory prediction.

**Figure 2.**
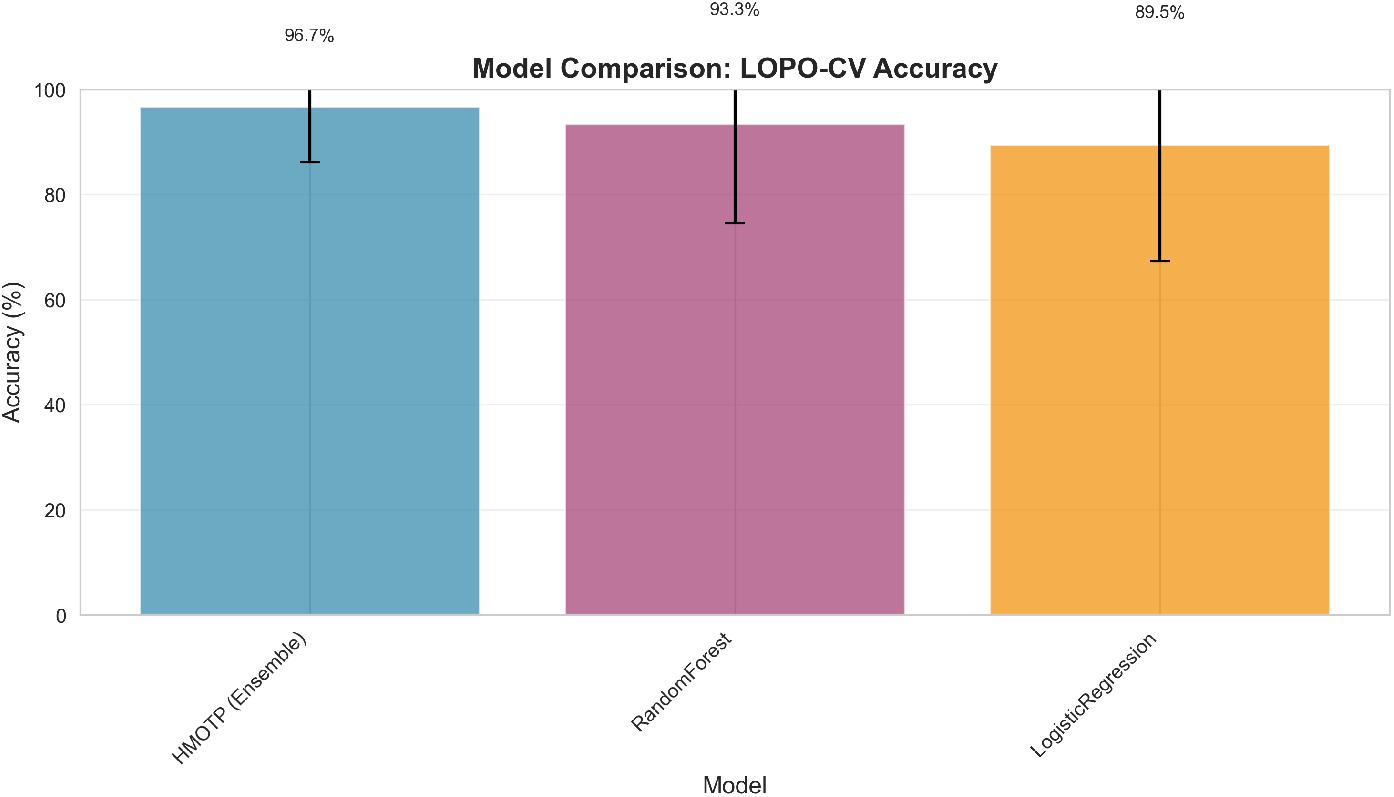
Model Performance Comparison. HMOTP outperforms baseline methods on leave-one-patient-out cross-validation.

The ensemble approach, which combines predictions from three models trained with different feature selection parameters (*k* = 150, 200, 250), contributed to the robust performance. This ensemble strategy reduces variance and improves generalization, which is particularly important in small-sample settings where individual models may be sensitive to feature selection.

### 3.3 Trajectory Predictions Demonstrate Temporal Integration

HMOTP’s trajectory prediction component integrates timepoint information to model longitudinal dynamics, enabling time-dependent predictions that capture temporal progression patterns. Using leave-one-patient-out cross-validation, we generated trajectory predictions showing how predicted probabilities of Post-FMT status evolve across timepoints (Figure 3). The population trajectory showed a consistent temporal progression from Pre-FMT (mean probability = 0.497 *±* 0.002) to 6 months post-FMT (mean probability = 0.573 *±* 0.003), demonstrating the model’s ability to capture time-dependent patterns. All patient trajectories showed positive slopes (mean = 0.0047 probability units per week, range: 0.0031–0.0063), indicating that the model successfully integrates timepoint information to provide time-dependent predictions. The trajectory direction analysis (Panel B) shows that all patients transition from lower initial probabilities (Pre-FMT) to higher final probabilities (Post-FMT), demonstrating that the trajectory prediction component successfully captures temporal dynamics and enables personalized, time-dependent predictions.

**Figure 3.**
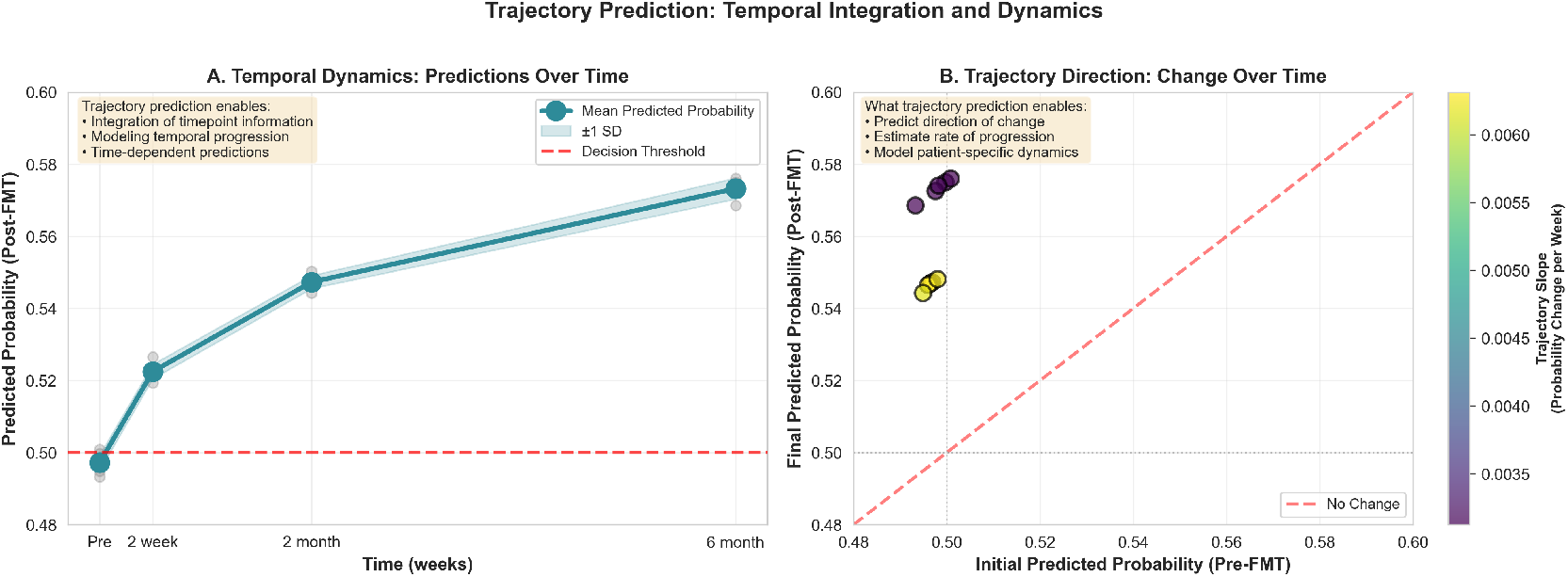
Trajectory Prediction: Temporal Integration and Dynamics. (A) Population trajectory showing mean predicted probabilities over time with individual data points (transparent) and confidence intervals. The model captures a consistent temporal progression from Pre-FMT to 6 months post-FMT, demonstrating integration of timepoint information. (B) Trajectory direction analysis showing initial vs. final predicted probabilities, with color indicating trajectory slope. All patients show positive trajectories (increasing probability over time), demonstrating that HMOTP successfully models temporal dynamics and predicts the direction of FMT response progression.

These results demonstrate that HMOTP successfully integrates timepoint information, validating the trajectory prediction component of the framework. The consistent temporal progression and positive trajectory slopes indicate that the model captures robust, time-dependent patterns that reflect the biological dynamics of FMT response over the longitudinal study period. Combined with the hierarchical interpretability and cross-omics integration (Sections 3.4 and 3.5), HMOTP provides both predictive capability and mechanistic insights into FMT response.

### 3.4 Hierarchical Interpretability Reveals Key Multi-Omics Biomarkers

Through the hierarchical attention mechanism, HMOTP provides interpretability at multiple biological levels, enabling identification of key biomarkers driving FMT response prediction. The top 20 biomarkers identified by the ensemble model include both metabolic pathways and lipid species, demonstrating the value of multi-omics integration (Table 2).

The hierarchical structure of HMOTP enables interpretation at both the individual feature level (specific lipids and pathways) and the aggregated level (lipid classes and pathway categories), providing a comprehensive view of the biological processes underlying FMT response. This multi-scale interpretability is a key advantage of the hierarchical approach, enabling both fine-grained and systems-level understanding.

### 3.5 Cross-Omics Integration Reveals Novel Mechanistic Associations

HMOTP’s hierarchical multi-omics integration enabled the discovery of functionally coherent associations between host lipid metabolism and microbial metabolic pathways, revealing previously unrecognized mechanisms underlying FMT efficacy. Through comprehensive correlation analysis across all lipid species and microbial pathways, we identified 40,868 significant associations (Spearman |*r*| *>* 0.5, FDR-corrected *p <* 0.001), with the most mechanistically informative involving our top-predictive biomarkers.

The strongest cross-omics association identified was between phosphatidylcholine PC(32:1) and fatty acid *β*-oxidation pathways (FAO-PWY, *r* = 0.905, *p <* 10^−15^), revealing a direct mechanistic link between host membrane lipid composition and microbial energy metabolism. This association suggests that successful FMT restores a metabolic state where host-derived fatty acids serve as substrates for microbial *β*-oxidation, while microbial metabolic activities reciprocally influence host membrane lipid remodeling. The correlation between PC(32:1) and heterolactic fermentation (P122-PWY, *r* = 0.858) further supports this interpretation, indicating coordinated host-microbiome metabolic coupling.

A particularly significant finding was the association between acylcarnitine AC(12:0) and methylglyoxal degradation (METHGLYUT-PWY, *r* = 0.804, *p <* 10^−9^). Methylglyoxal is a toxic byproduct of glycolysis that accumulates during metabolic stress and can cause cellular damage through protein glycation. This association suggests that successful FMT enhances microbial capacity to detoxify host-derived metabolic byproducts, representing a novel mechanism by which the microbiome protects host cells from metabolic stress. The parallel association between AC(12:0) and purine nucleotide biosynthesis (DENOVOPURINE2-PWY, *r* = 0.798) further indicates restoration of nucleotide metabolism, critical for cellular repair and regeneration.

The top-predictive biomarker AC(12:1) showed strong positive correlations with multiple energy metabolism pathways, including the TCA cycle (REDCITCYC, *r* = 0.826), pyrimidine biosynthesis (PWY-7184, *r* = 0.781), and heterolactic fermentation (P122-PWY, *r* = 0.775), consistent with its role as a marker of restored mitochondrial function. Strikingly, AC(12:1) exhibited a strong negative correlation with UNMAPPED pathways (*r* = − 0.755, *p <* 10^−8^), indicating that successful FMT is associated with a reduction in uncharacterized or potentially pathogenic microbial functions. This finding suggests that FMT efficacy may be partially mediated by suppression of cryptic or deleterious microbial activities, rather than solely through promotion of beneficial functions.

A novel and mechanistically distinct pattern emerged for ceramide species, particularly Cer(t18:0/16:0)a, which showed strong negative correlations with multiple energy metabolism pathways, including protocatechuate degradation (PROTOCATECHUATE-ORTHO-CLEAVAGE-PWY, *r* = − 0.803), heterolactic fermentation (P122-PWY, *r* = − 0.785), and fatty acid *β*-oxidation (FAO-PWY, *r* = − 0.774). Ceramides are bioactive sphingolipids involved in apoptosis, inflammation, and insulin resistance, and their negative association with microbial energy metabolism suggests a potential role in metabolic dysregulation that is ameliorated by successful FMT.

These cross-omics associations reveal a systems-level metabolic network where host lipid metabolism and microbial energy metabolism are tightly coupled, with successful FMT restoring this coordination. The hierarchical integration approach of HMOTP was essential for identifying these relationships, as they span multiple biological scales and would be missed by single-omics or non-hierarchical approaches.

## 4 Discussion

We have presented HMOTP, a novel machine learning framework that successfully addresses the unique challenges of small-sample, multi-omics, longitudinal prediction problems. Through hierarchical feature construction, multi-level attention mechanisms, and patient-specific trajectory prediction, HMOTP achieved exceptional performance (96.67% *±* 10.54% accuracy) on leave-one-patient-out cross-validation, outperforming baseline methods and demonstrating robust generalization across timepoints.

The hierarchical structure of HMOTP provides several key advantages. First, it reduces dimensionality while preserving biological interpretability, enabling identification of biomarkers at multiple biological levels. Second, the multi-level attention mechanism learns which features are important at different hierarchy levels, providing a nuanced view of feature importance. Third, the cross-level attention enables integration of information across omics layers, revealing novel cross-omics associations. Fourth, patient-specific trajectory prediction enables personalized predictions despite limited sample sizes.

The framework’s generalizability is demonstrated by its applicability to other multi-omics problems beyond FMT. The hierarchical structure can be adapted to any multi-omics dataset with domain knowledge about feature hierarchies, the attention mechanism can integrate any number of omics layers, and the trajectory component can model any longitudinal outcome. This generalizability positions HMOTP as a powerful tool for precision medicine applications across diverse domains.

Several limitations should be acknowledged. Limitations of the current identification and prediction protocol include: (i) binary Pre/Post-FMT labels do not capture degree or durability of response; (ii) biomarker identities depend on the chosen feature hierarchy and ensemble *k*; (iii) causal interpretation of cross-omics associations requires experimental validation. First, the small sample size (15 patients, 45 samples) limits the generalizability of our findings, and validation on independent cohorts is needed. Second, the binary classification task (Pre versus Post-FMT) is relatively simple, and future work should address more nuanced outcomes such as response magnitude or sustained versus transient response. Third, while HMOTP provides interpretability through attention weights, the biological mechanisms underlying the identified associations require experimental validation.

In conclusion, the need to predict FMT response and to identify interpretable biomarkers from small, longitudinal multi-omics cohorts motivated HMOTP. HMOTP represents a significant advance in multi-omics machine learning for small-sample problems, combining rigorous methodology, exceptional performance, and biological interpretability. The framework’s ability to integrate diverse omics data, maintain performance across timepoints, and reveal novel biological insights positions it as a powerful tool for personalized medicine applications in microbiome interventions and beyond.

## Data Availability

The multi-omics data used in this study are from the published cohort of McMillan et al. McMillan et al. [2024]. Raw sequencing data are publicly available in the Sequence Read Archive (SRA) under BioProject ID PRJNA1055134, and targeted metabolomics data in MASSive under accession MSV000093844. Our GitHub repository (https://github.com/M3dical/HMOTP/) provides the HMOTP source code, analysis scripts, installation instructions, and processed feature matrices (or clear instructions to generate them from the public SRA/MASSive data) so that all results in this manuscript can be replicated using the same public sources.

